# *FT*-like genes in Cannabis and hops: sex specific expression and copy-number variation may explain flowering time variation

**DOI:** 10.1101/2024.10.04.616617

**Authors:** Caroline A. Dowling, Todd P. Michael, Paul F. McCabe, Susanne Schilling, Rainer Melzer

## Abstract

*Cannabis sativa* is a fascinating, yet under-researched, species. To facilitate the global expansion of *C. sativa* cultivation a greater understanding of flowering time control is crucial. The *PEBP* gene family consists of universal promoters and repressors of flowering, with homologs of *FLOWERING LOCUS T* (*FT*) being highly conserved key regulators of flowering. *FT* encodes florigen, and balancing the florigen and anti-florigen signals is key for fine-tuning a crop’s flowering to local climatic conditions. Here, we provide an in-depth characterisation of the *PEBP* gene family in *C. sativa* and the closely related species *H. lupulus.* Phylogenetic analysis reveals expansion of *FT* and *TFL1/CEN* clades in the Cannabaceae. The retention of the duplicated *PEBP* genes likely has functional significance with divergent sequences and expression patterns hinting at signatures of sub-functionalisation. We speculate that duplicated *PEBP* genes have been crucial to the evolution of photoperiod insensitivity and sexually dimorphic flowering in *C. sativa* and harnessing the available genetic variation for these traits will be key for establishing this future crop.

## Background

Perhaps among one of the most infamous families of flowering plants is the Cannabaceae. This fascinating family includes *Cannabis sativa* (*C. sativa*) and *Humulus lupulus* (*H. lupulus*), both of which have cultural importance given the metabolites produced in glandular trichomes on female flowers [1,2]. Both species have tremendous commercial value, with *H. lupulus* used in brewing beer and uses of *C. sativa* spanning pharmaceuticals, biofuel and building materials [3]. However, the historic prohibition of *C. sativa*, has had the long-term impact of hindering in-depth research and breeding. While genetic studies on *C. sativa* remain in their infancy, the crop is quickly gaining attention [4–9].

Several interesting biological phenomena exist in the Cannabaceae, the genetic control of which can be uncovered via comparative genomic studies. Separated by 20 million years of evolution, *C. sativa* and *H. lupulus* are both dioecious but differ in their life history strategies with *C. sativa* being an annual and *H. lupulus* a perennial [10,11]. Furthermore, in both species, male individuals flower before females and the photoperiod strongly impacts plant development, with *C. sativa* and *H. lupulus* being among the first species for which the influence of photoperiod on flowering was demonstrated (Tournois 1912, cited in [12,13]). Both *C. sativa* and *H. lupulus* are short-day plants, but as *H. lupulus* is perennial, it requires long days to sufficiently develop vegetatively before transitioning to a reproductive state [10]. Therefore, understanding the genetic control of flowering, especially in the annual *C. sativa*, will be essential for expanding the geographic range of cultivation.

Since the 1930s, classical studies sought to uncover the underlying mechanism by which plants transition to a flowering state. The flowering stimulus was theorised to be a graft-transmissible flowering hormone, called florigen [14,15]. Eventually, extensive molecular genetic studies in the model plant *Arabidopsis thaliana* demonstrated that florigen is the protein product of *FLOWERING LOCUS T* (*FT*) [16]. *FT* is expressed in the leaf phloem companion cells and the mobile protein subsequently translocates to the shoot apical meristem where it interacts with transcription factors to initiate the floral transition [17].

*FT* belongs to the phosphatidylethanolamine-binding (*PEBP*) gene family. These genes have well-studied, diverse roles in plant growth and development, with the most prominent function being flowering time control [18]. There are three subclades of *PEBP*-like genes: *FT*-like, *TFL1/CEN*-like and *MFT*-like. Broadly speaking, *FT*-like genes are flowering promoters (florigens) whereas *TFL1/CEN*-like are flowering repressors (anti-florigens), and thus the balance between these opposing signals is a major determinant of reproductive development [18,19]. *MFT* has been phylogenetically demonstrated to be the sister subfamily to both, *FT* and *TFL1/CEN*-like genes, with a role in seed germination and dormancy [20,21]. However, functional diversification within these subclades has been reported, and amino acid substitutions at known positions can be indicative of whether a *PEBP* gene is a promoter or repressor of flowering [18,22].

*PEBP*-like genes have been targeted throughout domestication in many different species [23–27]. Previous phylogenetic studies of the *PEBP*-like gene family in other plant families like the Poaceae, Brassicaceae and Rosaceae [28–30] demonstrated that the copy number of *PEBP*-like genes can vary substantially. For example, within the Rosaceae, duplications of *PEBP* genes in the genome of *Malus domestica* have occurred likely due to an independent whole genome duplication unique to the *Malus* genus, resulting in eight *PEBP* genes [28]. Contrastingly, *Prunus persica*, which is also in the Rosaceae, possesses only five *PEBP* genes.

An angiosperm-wide analysis demonstrated that *FT* and *TFL1/CEN* clades have expanded differentially with a stark increase in copy number in monocots [31]. Increases in *PEBP* gene number are evident in *Sorghum bicolor* (19 genes), *Oryza sativa* (19 genes), and *Zea mays* (23 genes) [32–34]. Most monocots have five *FT* subclades, and preliminary analysis suggests sub-/neo-functionalisation within these lineages [31]. For example, in *Hordeum vulgare HvFT4* is involved not only in flowering but also in spikelet development [35].

Previous research has identified an apparent duplication of a *PEBP*-like gene, *CsFT1*, as a candidate gene contributing to photoperiod insensitivity (“autoflowering”) in *C. sativa* [8]. Furthermore, sexual dimorphism regarding flowering time in *C. sativa* means that investigating the *PEBP* gene family will likely yield fascinating insights [8]. To better understand flowering dynamics in *C. sativa* we characterised the *PEBP*-like gene family in the Cannabaceae. We provide a detailed overview of the *PEBP* gene family in *C. sativa* and *H. lupulus* and also explore *PEBP* diversity in newly released diverse *C. sativa* genomes. We find an unexpectedly large number of *PEBP* genes, specifically an expansion of both the florigen (*FT*) and anti-florigen (*TFL1/CEN*) clades in the Cannabaceae and hypothesise that functional diversification has occurred in this gene family in *C. sativa*. We hypothesise that *PEBP* gene duplications are part of the driving force behind photoperiod insensitivity and sexually dimorphic flowering in *C. sativa* and thus are key molecular breeding targets for the expansion of global *C. sativa* cultivation.

## Methods

### Identification of *PEBP*-like genes in *C. sativa* and *H. lupulus*

Annotated *PEBP* genes in *C. sativa* were obtained from the ‘CBDRx’ reference genome gene annotation (‘CBDRx’-cs10 v2; https://www.ncbi.nlm.nih.gov/assembly/GCF_900626175.2/, last accessed on 10 Jan. 2023) [36] and verified using BLAST with *A. thaliana* FT (AT1G65480) protein as query (e-value < 1e−5). Additionally, the ‘FINOLA’ genome (GCA_003417725.2) was searched for *PEPB*-like genes using NCBI BLAST with the ‘CBDRx’ PEBP genomic sequence as a query (e-value < 2e-68). The top hits were retained for each ‘CBDRx’ *PEBP* gene, checking the NCBI MSA viewer (v1.25.0) to ensure only full-length hits were retained.

For *C. sativa*genomes not on NCBI (https://resources.michael.salk.edu/resources/cannabis_genomes/index.html, last accessed on 18 Aug., 2023), BLAST databases were constructed using NCBI BLAST+ blastp (v2.10.1) (e-value < 1) with ‘CBDRx’ PEBP proteins as query: male genomes ‘Boone County’ (BCMa/BCMb), ‘Golden Redwood’ (GRMa/GRMb), ‘Ace High 3-2’ (AH3Ma/AH3Mb) and ‘Kompolti’ (KOMPa/KOMPb); female genomes ‘White Widow’ (WHWa/WHWb) and YunMa (YMv2a/YMv2b); monoecious genomes ‘Santhica 27’-2 (SAN2a/SAN2b) and KCDora (KCDv1a/KCDv1b) (Table S1, Figure S2). Nucleotide blast databases were generated for ‘Kompolti’ and ‘Boone County’ using NCBI BLAST+ makeblastdb to check for the presence of a previously identified helitron-like sequence in *CsFT1a* (QKVJ02000894.1|:191,793-192,866) [8].

For *H. lupulus* the most recent unmasked “Cascade” genome assembly and annotation [2] was downloaded from HopBase (http://hopbase.cqls.oregonstate.edu/dovetailDownloads/dovetailCascadeMasked.php, last accessed on 18 Aug., 2023). Dc-megablast (v2.10.1) was used with ‘CBDRx’ *PEBP* coding sequences as queries and an e-value cutoff of 0.001.

For genomes without a full annotation, (*H. lupulus* and *C. sativa ‘*FINOLA’) AUGUSTUS (http://bioinf.uni-greifswald.de/augustus/) and FGENESH+ (http://www.softberry.com/berry.phtml?topic=fgenes_plus&group=programs&subgroup=gfs, last accessed on 16 Aug. 2023) were used to predict protein sequence from blast hits [37,38]. The presence of a PEBP domain was verified using the NCBI conserved domain database [39]. Protein sequences were aligned with MAFFT and visualised in Jalview to check for an intact PEBP domain and the presence of a start codon [40,41]. For ‘FINOLA’ *PEBP*-like genes, any amino acid substitutions in comparison to ‘CBDRx’ were confirmed using the ‘FINOLA’ Illumina WGS short read data (SRS17330655, SRS17330649) that was generated previously [8].

To identify haplotigs (alleles assembled as two separate genetic loci), normalised coverage analysis was conducted. For *H. lupulus* publicly available Illumina WGS short read data (DRR024451) was obtained from NCBI SRA and mapped to the Dovetail “Cascade’’ reference genome (http://hopbase.cqls.oregonstate.edu/dovetailDownloads/dovetailCascadeMasked.ph <u>p</u>, last accessed on 8 May 2023) using BWA-MEM (Galaxy version v0.7.17.2) [42].

Coverage analysis was further conducted for ‘FINOLA’ using the aforementioned WGS short-read data [8] that were mapped to the ‘FINOLA’ genome (GCA_003417725.2, last accessed on 2 Jul., 2020) using BWA-MEM (Galaxy version v0.7.17.2) [42].

To get average coverage of blast hits, normalised relative to the whole genome coverage, samtools depth (-a, v1.10) and samtools bedcov (Galaxy Version 2.0.3) were used. An average coverage cutoff >0.8 was used. All megablast and coverage analyses were done using the Galaxy platform [43].

### Phylogenetic analysis of *PEBP*-like genes

For comparative sequence analysis of *PEBP*-like genes, a recent phylogenetic study was used to identify a Rosales species closely related to *Cannabis* with a high-quality reference genome, and *Prunus persica* was selected [44,45]. For *P*. *persica* gene identifiers were obtained from the literature [28], and sequences were obtained from NCBI. For *A. thaliana*, protein sequences were accessed from TAIR (The Arabidopsis Information Resource, www.arabidopsis.org, last accessed on 5 Sept., 2022). For *Glycine max* PEBP gene IDs were obtained from the literature [22,46], the newest gene annotation ID verified in Soybase (https://www.soybase.org/, last accessed 23 November 2023) and NCBI was used to acquire the newest RefSeq (Glycine_max_v4.0, GCF_000004515.6) protein sequence. Maximum likelihood phylogenetic trees were created as previously described [8]. In brief, protein sequences were aligned in MAFFT [41] and ALISTAT (v.1.3) masked residues with low completeness (> 0.5) in the alignment [47]. IQ-TREE was used for tree construction [48] with branch support values from Ultrafast bootstraps (UFBoot) and a Shi-modaira–Hasegawa approximate likelihood ratio test (SH-aLRT) test (1000 replicates each). A clade was deemed reliable if SH-aLRT and UFBoot supports were >80% and >95%, respectively. FigTree (v1.4.4, http://tree.bio.ed.ac.uk/software/figtree/, last accessed on 18 Aug. 2023) and ggtree [49] were used to visualise the trees.

### Synteny and gene structures analysis

Genomic sequences of various *C. sativa* PEBP were obtained and compared using dot plots generated with D-GENIES (https://dgenies.toulouse.inra.fr/, last accessed on 12 Dec. 2023). Synteny plots were generated using MCScanX and TBtools [50,51]. The MCScanX output was manually edited to ensure all verified HlPEBP hits were linked with their respective best CsPEBP blast hit in the synteny plot. For the ‘Kompolti’ synteny plots, MCScanX output was also edited to allow links between copies of duplicated genes. Publicly available gene annotation files (GFF3) were used for all genomes. The *H. lupulus* genome annotation was downloaded from HopBase (https://hopbase.org/content/cascadeDovetail/geneData/transdecoder/transdecoder <u>Output/transcripts.fasta.transdecoder.genomeCentric.gff3.gz</u>, last accessed 18 Dec., 2023). Transcripts corresponding to the *HlPEBP* genes were identified and if no transcript was found the GFF was manually edited (Table S2). For ‘FINOLA’ a GTF was constructed from RNA-seq data (see StringTie method). Gene structure plots accompanied by phylogenetic trees were visualised in TBtools, with MAFFT used to align protein sequences and IQtree to generate the tree [41,48].

### RNA-seq analysis

RNA-seq analysis was conducted as outlined previously [8,52]. Reads were mapped to ‘Kompolti’ using HISAT2 v2.2.1 [53]. The RSeQC Infer Experiment tool was used to determine the strandedness of mapped datasets [54]. StringTie was used to assemble transcripts and calculate gene expression in transcripts per million (TPM) [55]. Morpheus (https://software.broadinstitute.org/morpheus, last accessed on 13 November., 2023) was used to generate heatmaps from TPM values. RNA-seq data is available on NCBI from bioprojects PRJNA956491 and PRJNA1126191.

## Results

### PEBP genes identified in C. sativa and H. lupulus

In previous work, we identified 12 *PEBP*-like genes in *C. sativa* [8]. More specifically, we found four *FT*-like genes, four *TFL1/CEN*-like genes, and three *MFT*-like genes (Figure 1). To confirm the phylogenetic history and verify the number of the *C. sativa PEBP*-like genes identified, we analysed the *PEBP*-like genes in the closely related species *H. lupulus*.

**Figure 1.**
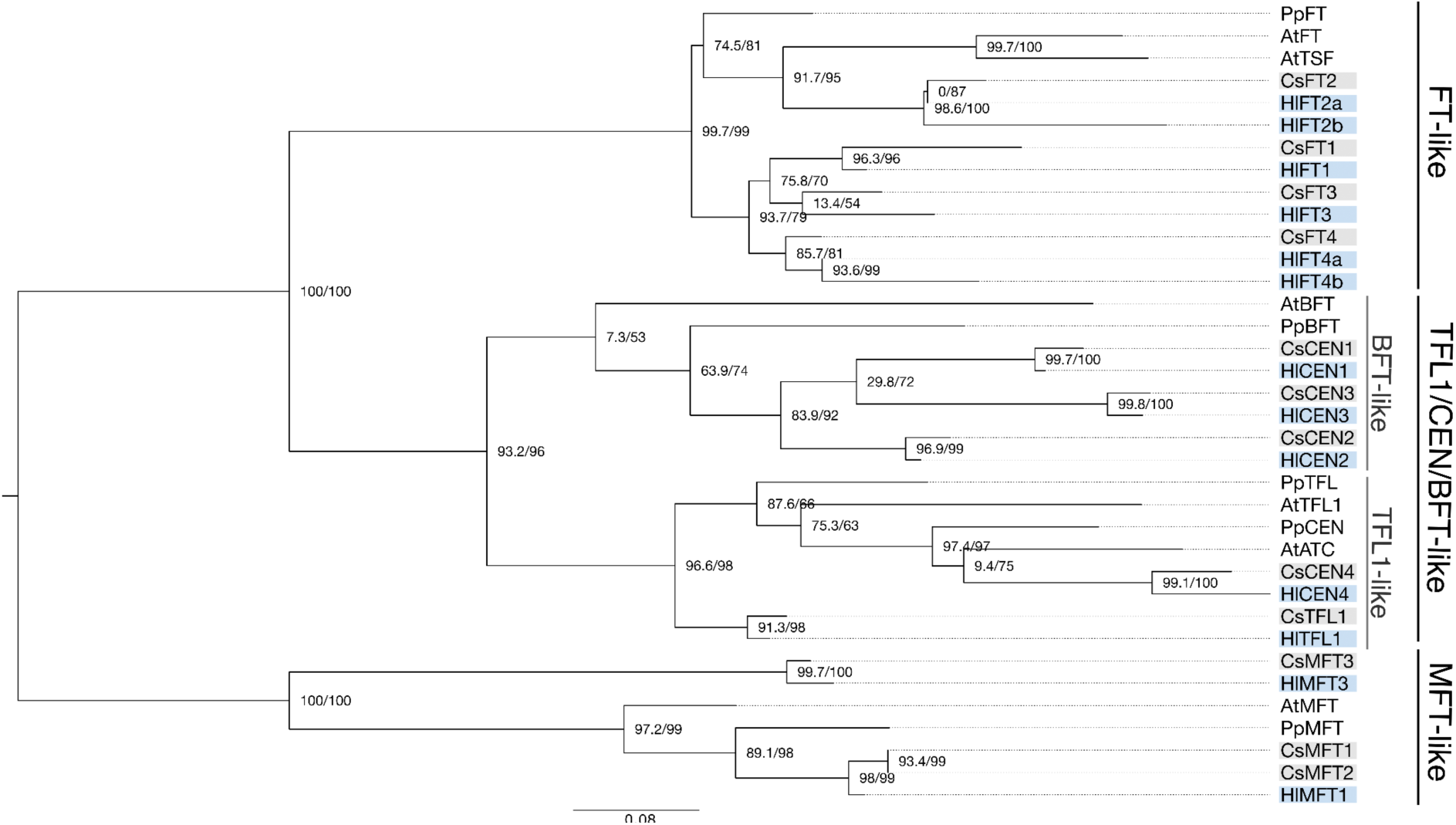
*PEBP*-like genes were identified in *C. sativa* and *H. lupulus.* A maximum likelihood phylogenetic tree of PEPB-like proteins from *A. thaliana, P. persica, C. sativa* and *H. lupulus* shows three main clades: FT, MFT and TFL1/CEN-like. Gene identifiers are outlined in Table S3.

To identify the *PEBP*-like genes in *H. lupulus*, dc-megablast was employed with 26 raw hits obtained when using the 12 *C. sativa PEBP*-like genes as query sequence (Table S2). Given the likelihood that “haplotigs” (haplotypes assembled as separate contigs) exist in the scaffold level *H. lupulus* assembly, the raw hits were subsequently verified using coverage and sequence analysis. After these filtering steps, 13 *PEBP*-like genes were verified to likely exist in *H. lupulus* (Figure 1, Table S2).

In our phylogenetic analysis, we also included *PEBP*-like genes from the model plant *A. thaliana* and from *Prunus persica* (peach). *Prunus persica* was chosen because like *C. sativa* it belongs to the Rosales [28,45].

The number of *TFL1/CEN/BFT*-like genes is the same in *C. sativa* and *H. lupulus* with five genes found in both species. A separation in a *BFT* and *TFL1/CEN* clade is visible (Figure 1). The *BFT* subclade follows, albeit with low support values, the species phylogeny: *A. thaliana BFT* is sister to all other *BFT*-like genes, followed by *P. persica*, *C. sativa* and *H. lupulus*. The three *BFT*-like genes found in *C. sativa* and *H. lupulus* show a 1-to-1 relationship. This indicates that *BFT*-like genes underwent two rounds of duplications in the lineage leading to *C. sativa* and *H. lupulus* after divergence from the *P. persica* lineage. In eudicots separate *TFL* and *CEN* clades can also be identified [31]. Though a *CEN* clade is resolved in our phylogeny (containing *AtATC, PpCEN, HlCEN4* and *CsCEN4*) a clear *TFL1* clade could not be reconstructed, although from an an evolutionary perspective it may appears plausible that *CsTFL1* is a *TFL1* ortholog.

There are two notable discrepancies in *PEBP* gene number between *H. lupulus* and *C. sativa*. Six *FT*-like genes were identified in *H. lupulus*, with apparent *HlFT2* and *HlFT4* duplications in comparison to *C. sativa* which has four *FT*-like genes (Figure 1, 2). The overall *FT* clade has strong support (99.7/99), while the internal separation into *FT2* and *FT1/3/4* subclades is sustained by lower support values (Figure 1). Within the *FT2* subclade are *CsFT2* and *HlFT2*, which are orthologous to *FT* from *A. thaliana*. The *FT1/3/4* subclade contains only *H. lupulus* and *C. sativa* genes and is not well supported (93.7/79, see support values in Methods) (Figure 1). The multiple *C. sativa* and *H. lupulus FT*-like genes found in this subclade indicate that multiple rounds of gene duplications took place. The *HlFT3* ortholog identified in the current *H. lupulus* assembly is a partial sequence and may represent a pseudogene.

**Figure 2.**
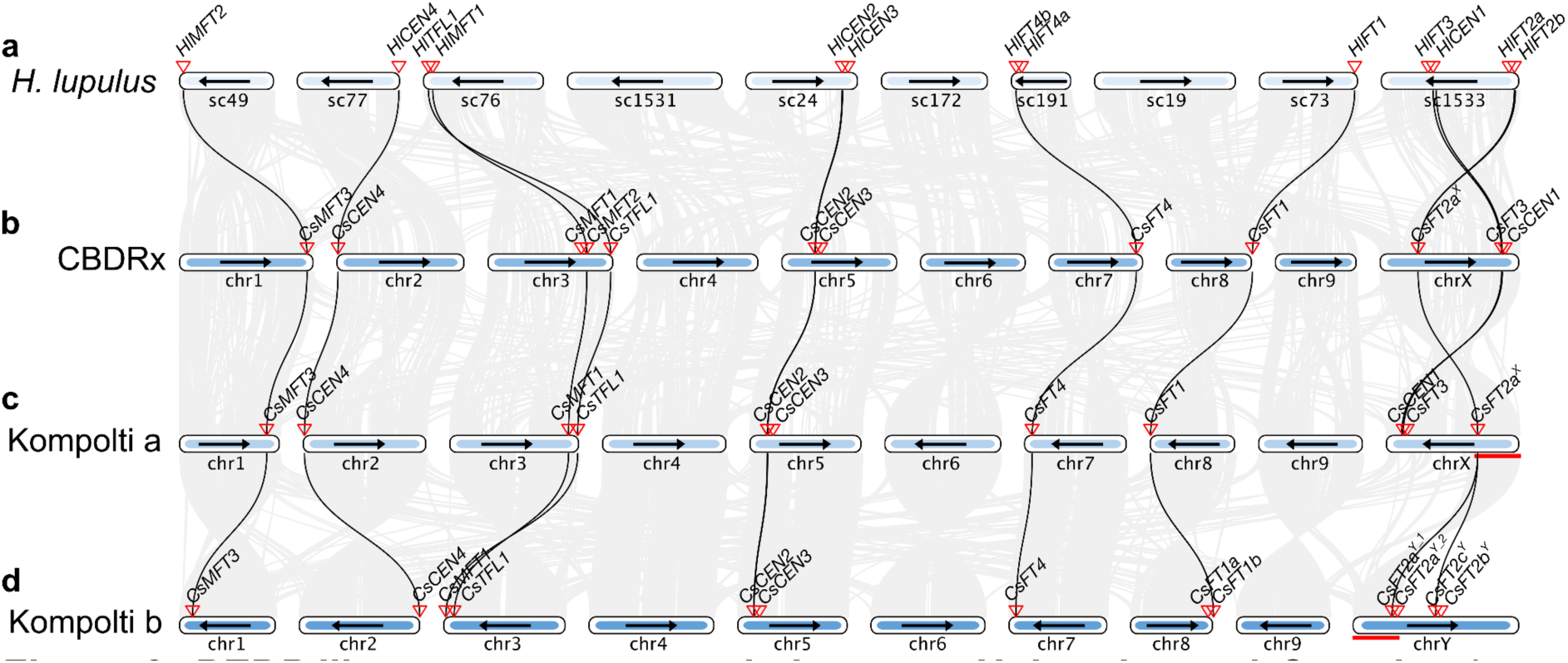
*PEBP*-like genes are syntenic between *H. lupulus* and *C. sativa.* Inter-genomic synteny blocks between the 10 largest scaffolds (sc) in *H. lupulus* (a) and the ten chromosomes of *C. sativa* accessions ‘CBDRx’ (female) (b) and ‘Kompolti’ (male). ‘Kompolti’ is a phased genome assembly, and thus both chromosomal haplotypes (c, d) were analysed. *H. lupulus* chromosomes are ordered as they relate to the *C. sativa* chromosomes [2]. Black arrows denote the chromosomal direction in a given assembly with respect to the ‘CBDRx’ genome. Inverted red triangles denote *PEBP* gene location. Grey lines between genomes denote synteny blocks from blast, with black lines denoting links between *PEBP* genes. Red lines denote the pseudoautosomal regions of the X and Y chromosomes of ‘Kompolti’.

Two and three *MFT*-like genes were identified in *H. lupulus* and *C. sativa*, respectively. Two separate *MFT* subclades exist in angiosperms [31], and this is supported in the present analysis. The *MFT1* subclade follows the species phylogeny: *A. thaliana MFT* is sister to all other *MFT*-like genes, followed by orthologs from *P. persica*, *C. sativa* and *H. lupulus*. The other *MFT* subclade contains *C. sativa CsMFT3* and *H. lupulus HlMFT2* only, which may indicate that the *CsMFT3* orthologs in *A. thaliana* and *P. persica* were lost (Figure 1). This is further supported when *Cucumis sativus* is included in the phylogeny, as this more distantly related species to *C. sativa* also has both *MFT* clades (Figure S1).

*CsMFT1* and *CsMFT2* in the ‘CBDRx’ *C. sativa* genome are almost identical in sequence and none of the other analysed *C. sativa* genomes displays this supposed duplication (see below), hence these paralogs might be haplotigs. To test for this, an additional coverage analysis was conducted for *CsMFT1* and *CsMFT2* using ‘FINOLA’ (which lacks the *CsMFT1-CsMFT2* duplication) and ‘Felina32’ (of unknown *CsMFT* genotype) short-read sequencing data mapped to ‘CBDRx’. In general, lower normalised coverage was found for *CsMFT1* and *CsMFT2* (0.3-0.7) than for *CsMFT3* (0.6-0.9) (Table S4). However, the results remain ambiguous as no short-read sequencing data is available for ‘CBDRx’. Therefore, the *CsMFT1* and *CsMFT2* duplication in ‘CBDRx’ may be genuine or an assembly artefact.

To understand the evolution of the *PEBP* gene family in the Cannabaceae, inter-genomic synteny analysis was conducted between *C. sativa* and *H. lupulus* (Figure 2a, 2b). A high level of macrosynteny exists in *C. sativa* and *H. lupulus*. Large interspecies collinear blocks exist for the *PEBP* genes, with all *PEBP* genes appearing in similar chromosomal locations in both species (Figure 2a, 2b). This analysis also indicated that the species-specific *H. lupulus* duplications observed in the phylogenetic tree (*HlFT2a/2b* and *HlFT4a/4b*) appear to be tandem duplications.

### *PEBP* copy number variation in other *C. sativa* genomes

Previously, we identified a duplication of *CsFT1* in the photoperiod-insensitive *C. sativa* accession ‘FINOLA’ [8]. Therefore, the *PEBP*-like genes in the publicly available ‘FINOLA’ genome [56] were analysed in greater detail. Using the same procedure as outlined above for *H. lupulus*, 15 *PEBP*-like genes were identified: Eight *FT*-like, four more than in the ‘CBDRx’ reference genome, four *TFL1/CEN*-like and two *MFT*-like genes (Table S5). The duplication of *CsFT1* in ‘FINOLA’ was previously reported [8]. The additional discrepancy in gene number is due to four *CsFT2* genes in ‘FINOLA’ compared to one in ‘CBDRx’.

To assess whether the *CsFT1* and *CsFT2* duplications found in ‘FINOLA’ are widespread in other *C. sativa* accessions, additional *C. sativa* genome assemblies that are fully phased (i.e. both chromosomal haplotypes are available) were analysed (http://salk-tm-pub.s3-website-us-west-2.amazonaws.com/cannabis_sativa/). Eight genomes were preliminarily analysed and were selected to try to encompass the diversity within the *C. sativa* gene pool (chemotype, flowering phenotype, sex, crop end-use) (Table S1, Figure S2). After preliminary phylogenetic analysis (Figure S2), a representative subset of four genomes was selected to supplement the in-depth sequence analysis on the grain cultivar ‘FINOLA’ and drug-type ‘CBDRx’: male hemp ‘Kompolti’, male feral hemp ‘Boone County’, a monoecious fibre-type ‘Santhica 27’ and a female drug-type ‘White Widow’.

The *CsFT1* duplication previously identified in ‘FINOLA’ was also detected in the accessions ‘Boone County’ and ‘Kompolti’ (Figure S3). In all analysed genomes, one *CsFT2* gene was detected, located on the sex chromosome X (*CsFT2a^X^*) (Figure 2b, 2c). In the male genomes ‘Boone County’ and ‘Kompolti’, three and four *CsFT2* duplicates were identified on chromosome Y, respectively (*CsFT2a^Y^, b^Y^, c^Y^*) (Figure 2d). *CsFT2a^Y^* is located at a similar position on the Y chromosome as CsFT2a^X^ on the X chromosome, within the pseudoautosomal region, but relatively close to the non-recombining region. Two *CsFT2a^Y^* genes located 0.3 Mb apart, *CsFT2a^Y_1^.K.b* and *CsFT2a^Y_2^.K.b*, were found on chromosome Y in ‘Kompolti’ (Figure 2d). The additional *CsFT2b^Y^* and *CsFT2c^Y^* genes were found central within the non-recombining region of chromosome Y in both ‘Kompolti’ and ‘Boone County’. For ‘FINOLA’ there is no well-supported chromosome location for *CsFT2*. The male individual, as well as the female individual, were both found to have a coverage of ∼1 for *CsFT2a*, while coverage for *CsFT2b* to *CsFT2d was* ∼0.5 in the male ‘FINOLA’ individual and 0 in the female ‘FINOLA’, indicating that those genes are located on the Y chromosome, similar to the other accessions with phased chromosome assemblies (Table S5).

Additionally, *CsCEN1* and *CsFT3* both appear to be only found on chromosome X (Figure 2b, 2c). In ‘FINOLA’, the coverage for these genes is ∼0.5 and ∼1-1.5 in male and female individuals respectively (Table S5), supporting the notion that they are located on the X chromosome. Accordingly, two alleles of *CsCEN1* and *CsFT3* (1 per chromosome X haplotype) were identified in female and monoecious genomes ‘Santhica 27’ and ‘White Widow’, while the genes are hemizygous in the male individuals of the accessions ‘Boone County’ and ‘Kompolti’ (Figure S3).

Furthermore, only two *MFT*-like genes are found in all genomes except ‘CBDRx’, consolidating that the putative *MFT* duplication in ‘CBDRx’ (*CsMFT1* and *CsMFT2*) is indeed cultivar-specific (Figure S3).

### Key amino acids denote likely PEBP function in *C. sativa*

PEPB-like proteins have characteristic amino acid residues determining their function in floral promotion or inhibition [18]. Five amino acids, in particular, are conserved in FT-like proteins compared to TFL1/CEN-like and MFT-like proteins (Figure 3): Y85, E109, Y134, W138 and Q140 [18,27,57,58]. A comprehensive list of PEPB-like proteins was compiled from the literature for comparison with *C. sativa* PEPB proteins (Table S4).

**Figure 3.**
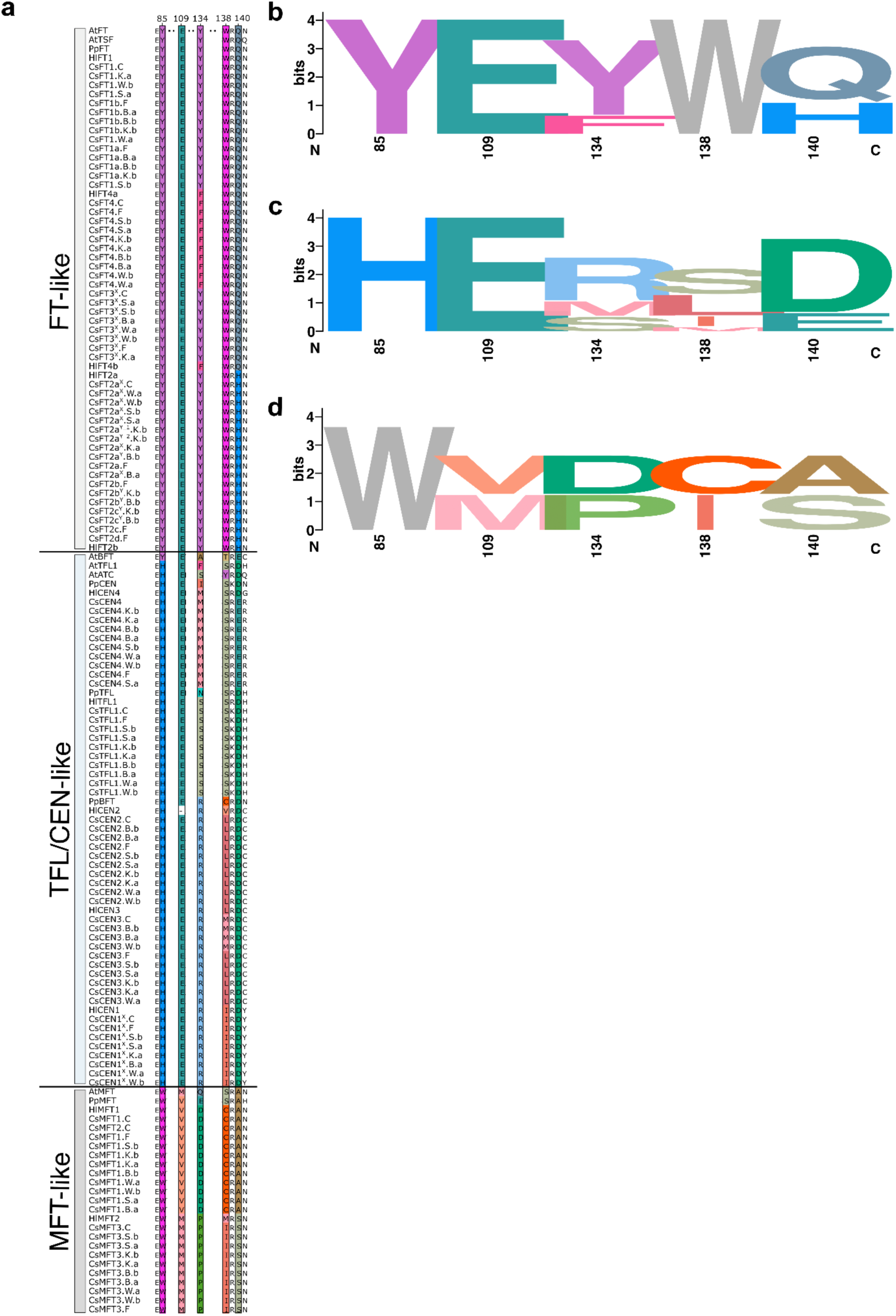
Key PEBP amino acid positions denote gene function. An alignment of PEPB-like proteins from *A. thaliana*, *P. persica*, *H. lupulus* and *C. sativa* shows three main clades: FT, MFT and TFL (a). Several amino acids are candidates for PEBP function (Table S3) but for concision, only five positions are highlighted here (position refers to AtFT): Y85, E109, Y134, W138 and Q140 [27]. Web logos summarise the variation at the key amino acids in *C. sativa* FT-like (b), TFL1/CEN-like (c), and MFT-like (d) proteins. HlFT3 was not included in (a) as it is only a partial sequence.

Out of these five key amino acid positions, three were identical between CsFT1 to CsFT4 and FT. At two positions changes compared to the canonical amino acids were observed: Y134F in CsFT4 and Q140H in CsFT2 (Figure 3). Y134F is also found in HlFT4 from *H. lupulus* and in the anti-florigen TFL1 from *A. thaliana*. Q140H is found in GmFT5a (from *Glycine max*) which phylogenetically clusters with CsFT2 and is a demonstrated flowering promoter (Figure S4) [59,60].

### Copy number variation of *CsFT1*

Previous research suggested that *CsFT1* duplicates in ‘FINOLA’ differed from one another in sequence (both protein and genomic) [8]. To investigate the variation at this locus between different *C. sativa* cultivars, the *CsFT1* sequences from a number of genomes were assessed in greater detail (Figure 4). In the phylogenetic tree and protein sequence alignment, three broad groups of *CsFT1* sequences exist: those similar to *CsFT1* (‘CBDRx’ and the photoperiod-sensitive cultivar ‘Felina 32’) [8], those similar to *CsFT1a* (photoperiod-insensitive cultivar ‘FINOLA’) and those similar to CsFT1b (‘FINOLA’) (Figure 4a). In total, eighteen amino acids vary when comparing all CsFT1 protein sequences (Figure 4a).

**Figure 4.**
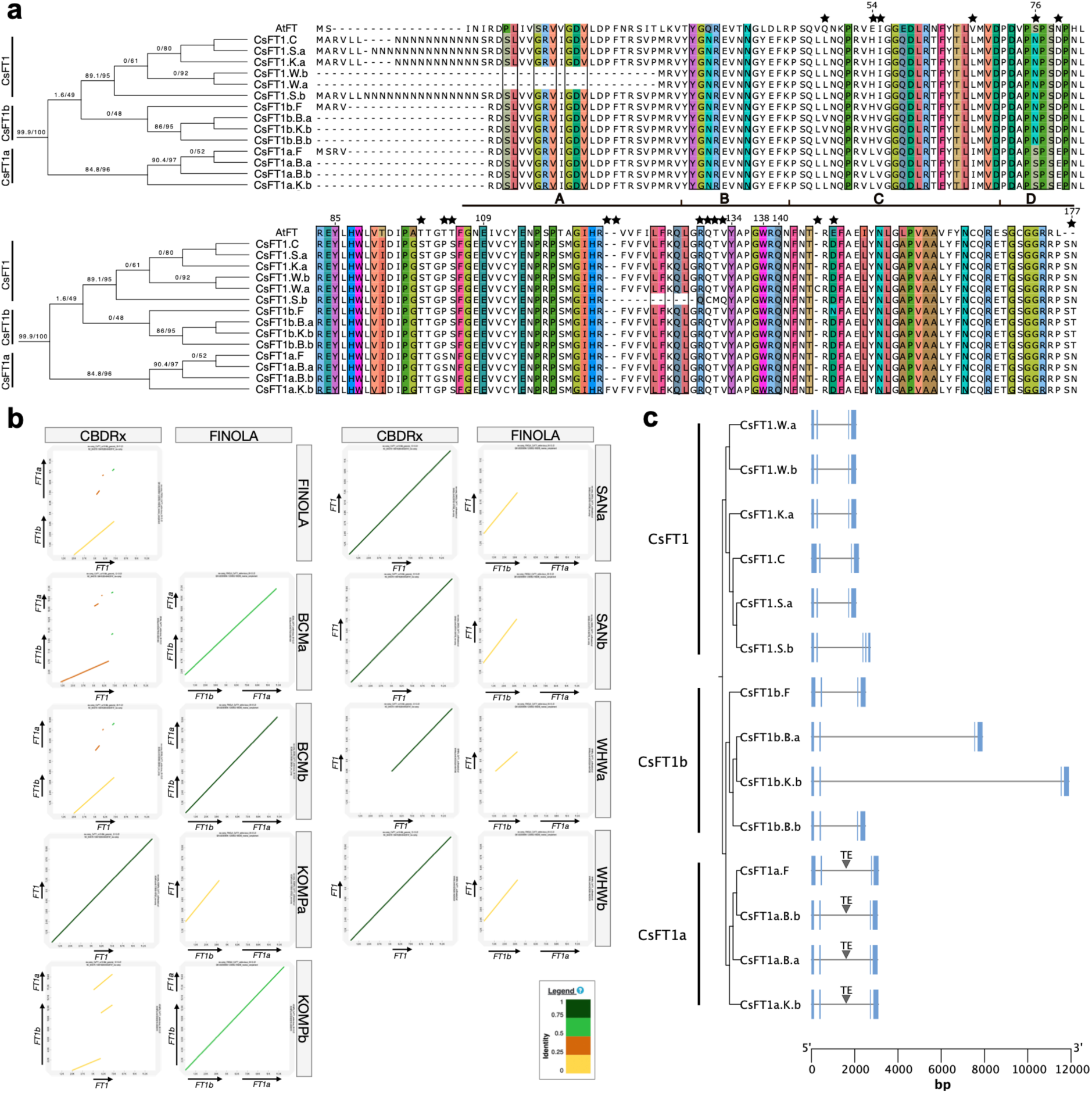
*CsFT1* copy number variation in various *C. sativa* genomes. An alignment of the CsFT1 protein sequences from the analysed *C. sativa* genomes and AtFT (a). Black stars denote amino acid differences between CsFT1 sequences. Numbered amino acid positions are with respect to AtFT. Highlighted AA columns are key for the function/designation of the PEBP subfamily (Table S3). The horizontal black line denotes the four segments encoded by the fourth exon [61]. The phylogenetic tree is according to Figure S3. Dot plots illustrating sequence similarity of *CsFT1* locus in various *C. sativa* genomes (b). Schematic of the *CsFT1* gene structure in various genomes, with sequences ordered according to the accompanying phylogenetic tree (c). Blue boxes denote the coding sequences. Gene labels include gene name, accession (C= ‘CBDRx’, F= ‘FINOLA’, W= ‘White Widow’, S= ‘Santhica 27’, K= ‘Komploti’, B= ‘Boone County’), and haplotype (a, b).

One of the amino acid differences between ‘CBDRx’ and ‘FINOLA’ CsFT1b is S76N, a position potentially important for function (Table S6). Three amino acid differences exist between all CsFT1a and CsFT1b alleles (amino acid position according to FT): H54L, N76S and N177T.

In ‘FINOLA’, ‘Kompolti’, ‘Boone County’ and ‘White Widow’ a CsFT1 protein start codon could not be predicted, potentially because the start codon is located on an upstream exon. In *CsFT1* of the ‘Santhica 27’ haplotype B, a 7 nucleotide insertion is predicted to result in an altered splicing pattern (Figure 4c), and a 10AA deletion in the protein sequence (Figure 4a).

Given that the protein sequences are relatively similar, the genomic *CsFT1* sequences were investigated. Dot plot comparisons of the ∼10-20 kb genomic fragments containing the *CsFT1* genes were analysed (Figure 4b). This confirmed that the locus covering *CsFT1* in ‘CBDRx’ vs. *CsFT1a* and *CsFT1b* in ‘FINOLA’ is quite diverged (Figure 4b). The dot plots further revealed that ‘FINOLA’, ‘Boone County’ and ‘Kompolti b’ all carry the *CsFT1a/1b* duplication. Similarly, the ‘CBDRx’ *CsFT1* locus resembles that of ‘Kompolti a’, ‘Santhica 27’ and ‘White Widow’. The genomic sequence upstream of the ‘White Widow a’ *CsFT1* gene is quite different from ‘CBDRx’ and ‘FINOLA’ and so does not appear on the dot plot (Figure 4b). However, these sequence deviations are upstream of the *CsFT1* coding sequence (Figure 4b).

Intronic variation was observed in the *CsFT1* duplicates with a transposable element found in the large intron of *CsFT1a* [8]. The same helitron-like sequence was found in *CsFT1a* of ‘Boone County’ and ‘Kompolti’ (Figure 4c). The intron length of *CsFT1a*-like alleles is equal across accessions, whereas the large intron of *CsFT1b* in ‘Boone County’ and ‘Kompolti’ varies in length (Figure 4c).

### Copy number variation in *CsFT2* in male *C. sativa*

All CsFT2 protein sequences appear to be very similar, with only five amino acids differing between the various CsFT2 sequences (Figure 5a).

**Figure 5.**
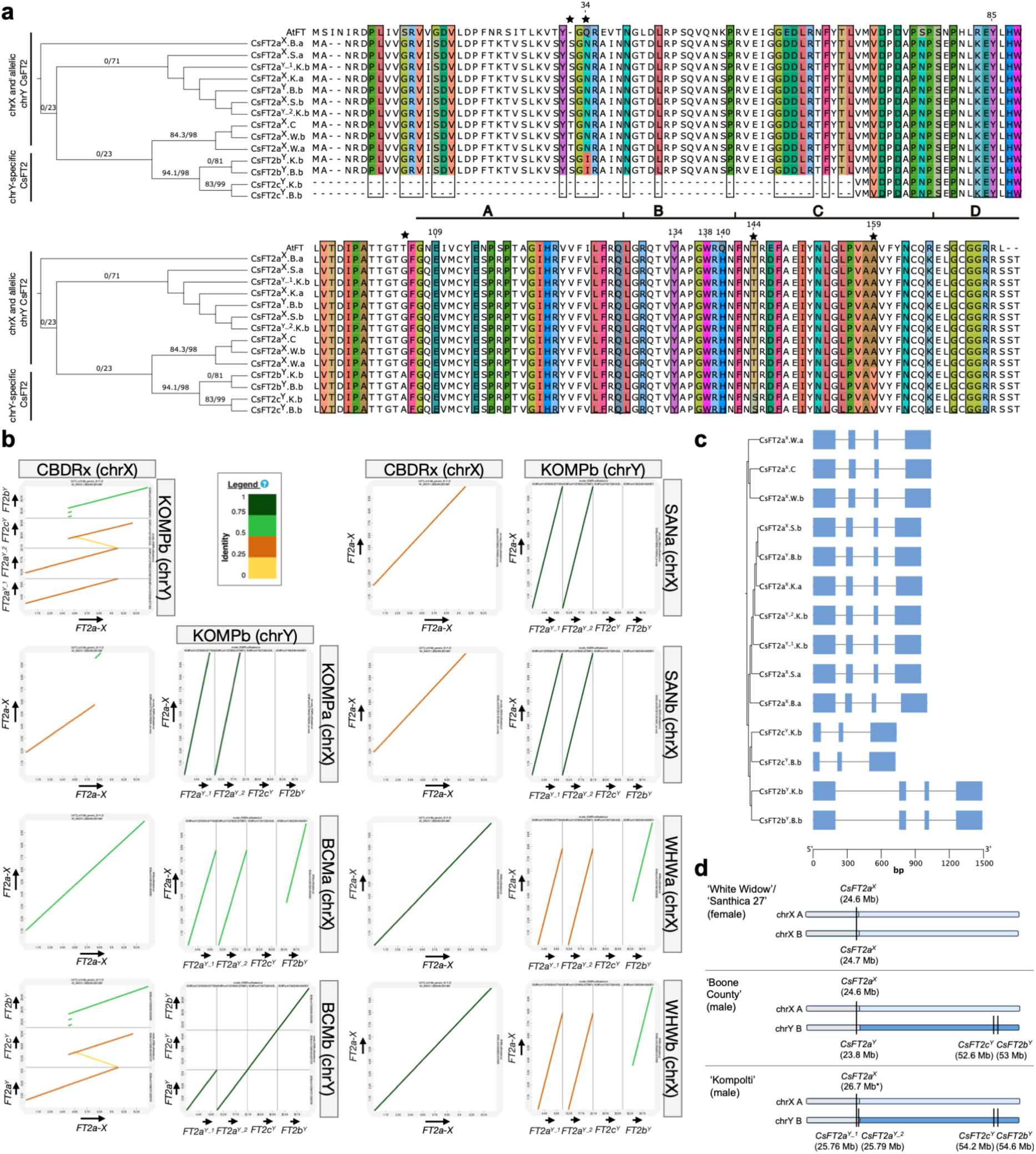
*CsFT2* copy number variation in various *C. sativa* genomes. (A) An alignment of the CsFT2 protein sequences from the analysed *C. sativa* genomes and AtFT. Black stars denote amino acid differences between CsFT2 alleles. Highlighted AA columns are key for the function/designation of the PEBP subfamily (Table S3). The horizontal black line denotes the fourth exon of the protein, split into four segments [61]. The phylogenetic tree is cropped from Figure 3. (B) Dot plots illustrating the genomic landscape of the *CsFT2* locus in various genomes. (C) Schematic of the *CsFT2* gene structure in various genomes analysed, with sequences ordered according to the accompanying phylogenetic tree. Blue boxes denote the coding sequences. Gene labels include gene name (-X or -Y if located on sex chromosomes), accession (C= ‘CBDRx, F= ‘FINOLA’, W= ‘White Widow’, S= ‘Santhica 27’, K= ‘Komploti’, B= ‘Boone County’), and haplotype (a, b). (D) Schematics demonstrating the allele combinations present in female (‘White Widow’/’Santhica 27’) and male (‘Boone County’/‘Kompolti’) genomes. Grey, light blue and dark blue boxes denote the pseudoautosomal, chrX-specific and chrY-specific chromosomal regions respectively. Vertical black lines pinpoint the genomic location of *CsFT2* alleles. *Expressed the genomic location of Kompolti chrX CsFT2a to ensure the same chromosomal direction as other assemblies.

Three of those amino acids differences are at positions potentially important for function between CsFT2a and CsFT2b^Y^/CsFT2c^Y^ (Figure 5a, Table S4, amino acid position according to FT): N34I in CsFT2b^Y^, T144S in CsFT2c^Y^ and A159V in both CsFT2b^Y^ and CsFT2c^Y^. Furthermore, CsFT2c^Y^ found in both ‘Boone County’ and ‘Kompolti’ appears to be shorter at the N-terminal end (Figure 5a, b, c).

Dot plot analysis was conducted to compare the genomic fragments containing *CsFT2* duplicates (Figure 5b). ‘FINOLA’ was not included in the in-depth *CsFT2* analysis as it lacks a chromosome Y assembly, and thus ‘Kompolti’ was used for genomic comparison.

In chromosome X haplotypes (‘CBDRx’, ‘Kompolti A’, ‘Boone County A’, ‘Santhica 27’ and ‘White Widow’) only *CsFT2a^X^*was present (Figure 5b). In chromosome Y haplotypes (‘Kompolti B’ and ‘Boone County B’) three *CsFT2a^Y^ to c^Y^* were found. In ‘Boone County’ and ‘Kompolti’ CsFT2c^Y^ is shorter than the ‘CBDRx’ query and has sequence similarity to ‘CBDRx’ in both the forward and reverse directions, resembling a pseudo-palindromic sequence (Figure 5a, b). Overall the dot plots provide a complex picture, with different levels of similarity between *CsFT2* loci. For example, the *CsFT2a^X^* locus appears to be very similar between ‘CBDRx’ and ‘White Widow A’, but different between ‘CBDRx’ and ‘Kompolti A’ (Figure 5 b).

The gene structures of *CsFT2* alleles were visualised, and the largest variation was observed in the introns of *CsFT2b^Y^* and the lack of exon 1 in *CsFT2c^Y^*(Figure 5c).

In summary, the *CsFT2* gene number appears to be dynamic on Y chromosomes in *C. sativa* (Figure 5).

### Expression analysis of *PEBP* genes at vegetative, floral transition and flowering suggests functional conservation

To assess whether the identified genes are expressed and potentially functional during the floral transition, the expression of all *PEBP*-like genes in *C. sativa* was analysed in vegetative and flowering tissues from the dioecious photoperiod insensitive cultivar ‘FINOLA’, the monoecious photoperiod sensitive cultivar ‘Felina 32’, and F_2_ individuals from a ‘Felina 32’ × ‘FINOLA’ cross described previously (Figure 6) [8].

**Figure 6.**
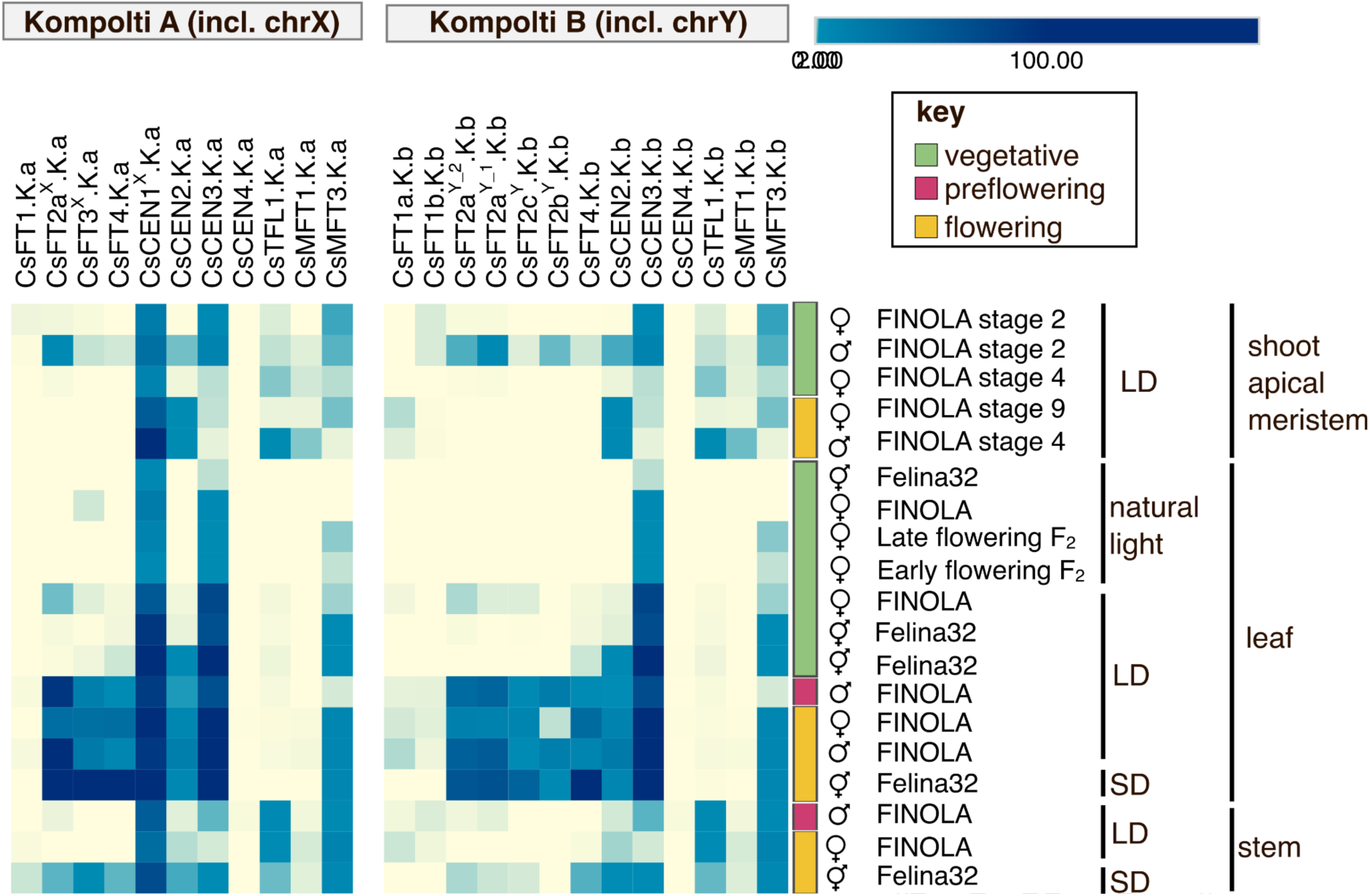
*PEBP*-like gene expression in *C. sativa.* Heatmap of RNA-seq data performed on flowering and vegetative tissue samples from ‘FINOLA’, ‘Felina 32’ and F_2_ individuals, mapped to ‘Kompolti’ haplotype A and B. Each row is an average TPM of the biological replicates. Each sample type has 3-5 biological replicates. Growth stages refer to the number of true leaf pairs [52]. Note that, although all transcripts were mapped to both Kompolti haplotypes, the female and monoecious plants have no Y chromosome. Transcripts from female or monoecious plants mapping to Y chromosomal genes probably originated from X chromosomal *FT* genes. Likewise, ‘Felina 32’ encodes *CsFT1* and ‘FINOLA’ encodes both *CsFT1a* and *CsFT1b*.

The male ‘Kompolti’ genome was used as a reference for RNA-seq analysis and as ‘Kompolti’ is a phased assembly, both haplotypes are available. ‘Kompolti A’ includes one *CsFT2* gene on chromosome X and one *CsFT1* gene on chromosome 8, whereas ‘Kompolti B’ possesses the chromosome Y *CsFT2* genes and the *CsFT1* gene duplication on chromosome 8 (Figure 2c, 2d, 6). This allowed the expression of the duplicated *PEBP*-like genes to be visualised using an appropriate reference genome.

As outlined previously [8], in ‘Felina32’ *CsFT1* expression was only observed at flowering under short-day conditions (Figure 6), whereas in ‘FINOLA’ either *CsFT1a* or *CsFT1b* expression was found in all developmental stages and all tissue types (Figure 6b), but primarily in flowering samples with *CsFT1a* being higher expressed than *CsFT1b* (Figure 6).

*CsFT2a* is highest expressed in leaves at flowering, though some expression is found in vegetative leaves and shoot apical meristem and flowering stem (Figure 6). *CsFT2b* and *CsFT2c* have similar expression patterns, mostly restricted to male and female leaves at flowering and male vegetative shoot apical meristem (Figure 6).

The expression patterns of *CsFT3* and *CsFT4* are similar and highest in leaf samples at flowering. In ‘Felina32’ both *CsFT3* and *CsFT4* expression were detected in stems at flowering, but the expression was absent in ‘FINOLA’ samples at the same stage (Figure 6a).

Expression of *CsTFL1* is highest in stems and the shoot apical meristem, with only trace expression found in leaf tissues (Figure 6a, b). *CsCEN1* is highly expressed in all samples (Figure 6a). *CsCEN2* is highest at flowering with expression in leaves and the shoot apical meristem (Figure 6a, b). *CsCEN3* is expressed in all tissues and is highest in leaves both in vegetative samples and at flowering (Figure 6a, b). *CsCEN4* expression was not detected in any samples (Figure 6a, b). *CsMFT3* is higher expressed than *CsMFT1* in all samples, with only trace expression found of the latter in shoot apical meristem and stem (Figure 6a, b).

## Discussion

### Clade-specific expansion of the *PEBP* gene family in the Cannabaceae

*PEBP* genes have crucial roles in developmental processes, especially the floral transition [18]. Previous research identified *PEBP*-like genes as candidates for flowering in *C. sativa* [8], and the growing availability of genomes of various accessions allowed an exhaustive search of this gene family to be performed in *C. sativa* and the closely related species *H. lupulus*.

Our phylogenetic analysis reveals that in *C. sativa* and *H. lupulus,* both the *FT* and *BFT* clades have expanded compared to *P. persica* and *A. thaliana* (Figure 1). The expansion is most obvious in the *FT* clade. The topology of the *FT* clade is likely not fully resolved given the low support values for some of the branches. The current topology would support gene losses in the lineages leading to *A. thaliana* and *P. persica*. Instead, we consider it more likely that there were several rounds of *FT* duplications in the lineage leading to the Cannabaceae.

For most *PEBP* genes in *C. sativa* and *H. lupulus*, a 1-to-1 orthology is observed, indicating that most gene duplications occurred before *H. lupulus* and *C. sativa* separated. An exception is *HlFT2* and *HlFT4* which appear to be duplicated specifically in *H. lupulus*.

It is interesting to note the location of *PEBP* genes in common syntenic blocks between *C. sativa* and *H. lupulus* genomes (Figure 2). This further supports that the expansion of the *PEBP* gene family within the Cannabaceae occurred before the speciation event leading to *C. sativa* and *H. lupulus* 25 million years ago [3,62], but after the separation of the Cannabacae within the Rosales, given that *P. persica* of the Rosaceae possesses only five *PEBP*-like genes (Figure 1, 2).

The origin of the duplications in the *FT* and *BFT* clades is unclear. Many increases in *PEBP* gene number in the eudicots are whole genome duplication (WGD)-mediated (*Malus domestica*, *Glycine max*) [22,33]. However, the recent *C. sativa* pangenome study suggests that *C. sativa* has not had a recent WGD event [63].

Small-scale duplications can also contribute to copy number variations, though it can be difficult to ensure a small-scale duplication is not an undetected remnant of an ancient whole genome duplication event [64,65]. Small-scale duplications can arise in several ways (tandem, segmental, transposon-mediated, retroduplication) and an accurate estimation of the extent of these occurrences can be limited by the quality of genome assemblies such as paralogues assembled as a single gene [65]. Investigating the potential origins of the *PEBP* duplications in the Cannabaceae will be interesting as more closely related species gain high-quality genome assemblies.

Nonetheless, the functional consequences of retaining so many duplicated *PEBP* genes are intriguing.

### Sequence divergence in duplicated *PEBP* genes indicates functional divergence

The retention of an expanded *PEBP* gene family in the Cannabaceae suggests that the duplicated *PEBP*s may have adaptive significance.

For all identified *C. sativa* and *H. lupulus PEBP*-like genes the key amino acids denoting flowering promoter/repressor function were analysed (Table S6). In the majority of cases, the phylogenetic location of the analysed *PEBP* genes matches the expected amino acid residues that indicate flowering-promoting or repressive activity (Figure 3, Table S6). Sequence divergence may be indicative of functional divergence, and in two cases sequence divergence in key amino acids was observed: CsFT2 and CsFT4 (Figure 3). In several species such as *O. sativa*, *Populus* spp., and *G. max* duplicated *PEBP* genes have undergone sub- and neo-functionalisation to regulate flowering as well as other developmental processes under particular environmental conditions [18,66].

CsFT2 has four out of the five residues expected to be present in a floral promoter, the exception being Q140H (Figure 3). The *G.max* protein GmFT5a which is orthologous to CsFT2 also has the same substitution (Figure 5, S3). Intriguingly, *GmFT5a* has a demonstrated role for latitudinal adaptation in *G. max*, and drives flowering in long-day conditions [27,60]. Given the phylogenetic relationship between GmFT5a and CsFT2, this could indicate that CsFT2 may have a similar role in photoperiodic flowering in *C. sativa*.

The male-specific duplicates of *CsFT2, CsFT2b^Y^* and *CsFT2c^Y^*, could be responsible for sexual dimorphic flowering in *C. sativa* given the expression of these genes just before and at flowering (Figure 6). In CsFT4 and HlFT4, Y134F substitution is observed compared to the other FT-like proteins (Figure 3). Tyrosine (Y) and phenylalanine (F) are similar amino acids, and so one can speculate that this has only modest implications for the function of the proteins. However, the floral repressor AtTFL1 also has phenylalanine at this position, as does HvFT4, a change-of-function FT that represses flowering in *H. vulgare* (Figure 3) [35]. Conversely, PsFTc, ZCN8 and HvFT3 also have Y134F and are flowering promoters in *Pisum sativum*, *Zea mays* and *H. vulgare*, respectively [67–70]. Therefore it remains to be seen whether CsFT4 and HlFT4 are floral repressors in the Cannabaceae. Interestingly, *CsFT4* is expressed in vegetative leaves under long-day conditions in the photoperiod-sensitive cultivar ‘Felina 32’ which may indicate anti-florigen activity (Figure 6).

*CsFT3* and *CsFT4* seem to be co-expressed in most sampled tissues which may indicate redundant or complementary roles in promoting flowering. The best hit for *FT3* in *H. lupulus* is only a partial sequence (Figure 1, 2).

### *PEBP* expression patterns indicate conservation as well as functional divergence

A divergence in the expected expression pattern from comparisons to other species may be indicative of a divergence in function within the PEBP subclades. In *C. sativa* and *H. lupulus* both angiosperm *MFT* clades are retained whereas one *MFT* is lost in *P. persica* and *A. thaliana* (Figure 1). The more distantly related species *Cucumis sativus* also possess two MFT clades, suggesting that the two clades found in the Cannabaceae are indeed the ancestral angiosperm duplication of MFT (Figure S1) [31]. Genes of the *MFT* clade typically function in seed dormancy [18,20,21]. The differing expression patterns observed in *CsMFT3* and *CsMFT1* may indicate that the genes have sub-functionalised (Figure 6). *CsMFT3* is expressed in all analysed tissue types and at all stages, whereas *CsMFT1* expression is restricted to shoot apical meristem and stems (Figure 6). In *O. sativa MFT* homologs have diverged in function as well as in expression pattern. *OsMFT1* is highly expressed in various tissues including leaves, seeds and panicles and functions in promoting panicle development and drought tolerance but delays heading date [71,72]. In contrast, *OsMFT2* expression is more specific to seeds and is a negative regulator of seed germination and positive regulator of seed dormancy, functions shared with *MFT* orthologs in *Triticum aestivum* and *A. thaliana* [20,21,73]. Therefore, it may be the case that in *C. sativa*, *CsMFT1* has a conserved function in seed development whereas the function of *CsMFT3* has similarly diversified as *OsMFT1* has in *O. sativa* and may operate in abiotic stress adaptation and flower development as well as flowering.

Differential expression patterns in the *C. sativa CEN/TFL1* clade suggest non-redundant functions in repressing flowering. The spatial pattern of *CsTFL1* expression suggests a conserved function in determining plant architecture as in other species, given the higher expression observed in shoot apical meristem and stems [18,74]. No expression was detected for *CsCEN4* which could indicate that this is a pseudogene or expressed under specific conditions not analysed. The remaining three *CsCEN*-like genes are phylogenetically closer to one another, albeit with low branch support. *CsCEN1*, though located on chromosome X, is highly expressed in males and females, in all tissues and at all developmental stages (Figure 6). *CsCEN2* expression mainly occurs in pre-flowering and flowering individuals, while *CsCEN3* expression is found in both vegetative and flowering samples, suggesting an earlier function in development. Overall, these expression patterns suggest that *CsCEN1, CsCEN2* and *CsCEN3* may have differential but important roles in repressing flowering at different developmental stages and in different tissues. In *A. thaliana BFT* is associated with delayed flowering under high salinity, while in *Populus* spp. and *Actinidia chinensis* (kiwifruit), *CEN/TFL1*-like genes regulate seasonal flowering and dormancy release [75–77]. Thus, members of the *CEN/TFL1* clade may be interesting candidates for investigating the differences in flowering regulation in the annual *C. sativa* and perennial *H. lupulus* in the Cannabaceae.

### Sex chromosome gene duplications may explain sexually dimorphic flowering in *C. sativa*

Sexual dimorphic flowering time has been previously reported in *C. sativa* [8,78,79] and *H. lupulus* [80,81]. However, the genetic underpinnings conferring flowering sexual dimorphism are unknown. In previous research, we analysed a *C. sativa* population segregating for two loci associated with photoperiod insensitive flowering, *Autoflower1* and *Autoflower2* [8]. When cultivated in non-inductive long-day conditions, some individuals that did not carry the photoperiod insensitivity conferring alleles of *Autoflower1* and *Autoflower2* still flowered. All of those plants were male. This suggested the existence of a third photoperiod related locus on the *C. sativa* Y chromosome. We speculate that the earlier flowering observed in *C. sativa* males may be associated with *PEBP* genes on the sex chromosomes, namely the chromosome Y-specific *CsFT2* duplicates (Figure S4, 6).

On chromosome Y several *CsFT2* copies are found, in ‘Kompolti’ termed *CsFT2a^Y_1^, CsFT2a^Y_2^, CsFT2b^Y^, CsFT2c^Y^* (Figure 2, 3, 5d). The exact number of *CsFT2* duplicates on chromosome Y may be overrepresented in the current assemblies, but at least two full-length *CsFT2* copies are likely located on chromosome Y (Figure 5). *HlFT2* is also located on chromosome X in *H. lupulus.* It would be interesting to delve into the evolutionary history of these duplicates and whether they also precede the speciation event 20 million years ago leading to *C. sativa* and *H. lupulus* [82], but uncovering this would require a chromosome Y assembly in *H. lupulus* which is currently lacking.

As aforementioned, *HlFT2* an ortholog of *CsFT2* is also located on chromosome X in *H. lupulus.* A recent genome assembly of *Ficus hispida* of the Moraceae family (one of the closest related families to the Cannabaceae and predominantly dioecious) also identified male-specific *FT*-like genes [83]. *FT* orthologs have also been detected in the male-specific region in *Amaranthus tuberculatus* and *Amaranthus palmeri* where it is postulated to contribute to male fitness [84,85]. In *Ambrosia artemisiifolia*, earlier male flowering increases reproductive success [86]. Therefore it will be interesting to assess the connection between reproductive success and male flowering variation in *C. sativa*, in addition to investigating the evolutionary history of *FT* duplications on sex chromosomes in species closely related to *C. sativa*.

Uncovering the genetic control of sexually dimorphic flowering in *C. sativa* would have several conceivable applications. Future breeding efforts may seek to substantially delay male flowering until the cannabinoid-producing female flowers have matured and are ready for harvest. Alternatively, it is more conceivable to have extremely early flowering males that would complete their life cycle faster and thus wouldn’t compete with females for resources (light, space, nutrients) in the field. Finally, it may be a target to synchronise male and female flowering for fibre hemp, as the lignification that coincides with flowering reduces fibre quality [87,88], and thus synchronising the flowering time of both sexes would ensure the largest possible yield, with the highest fibre quality.

### Photoperiod insensitivity in *C. sativa* may be a stepwise system involving *CsFT1* duplications

Previous research has demonstrated a duplication of *CsFT1* in a QTL for the photoperiod-insensitive flowering in ‘FINOLA’ [8]. In this study, we showed that this duplication also exists in two publicly available genomes ‘Kompolti’ and ‘Boone County’ (Figure 2, 3). At the protein sequence level, there are three main groups of alleles: those resembling *CsFT1* (‘CBDRx’-like), those resembling *CsFT1a* (‘FINOLA’) or *CsFT1b* (also ‘FINOLA’) (Figure 4a). Interestingly, the sequenced ‘Kompolti’ individual is heterozygous for the duplication and thus possesses all three allele types (Figure 4a, b, c). ‘Boone County’ is homozygous for the duplication, *CsFT1a* and *CsFT1b* (Figure 4a, b, c).

Although several substitutions between different CsFT1 sequences were identified, only one amino acid of potential functional importance was clade-specific: N76S. This substitution differentiates CsFT1a (S76) from CsFT1 and CsFT1b (N76) allele types (Figure 4a). In *A. thaliana* S76 may be important for the efficient accumulation of FT in the shoot apical meristem, though the substitution tested in that study was S76A [89]. Thus the impact of S76N in ‘CBDRx’-like CsFT1 and CsFT1b is unknown, though it is interesting that CsFT1a has S76, the same amino acid as *A. thaliana* at this position (Figure 4a). The presence of serine at position 76 in CsFT1a may facilitate more efficient unloading of FT protein from the phloem into the shoot apical meristem, contributing to photoperiod-insensitive flowering.

At the genomic level, dot plots confirmed the similarity between the ‘FINOLA’, ‘Boone County’ and ‘Kompolti’-B *CsFT1* duplication (Figure 4b). We previously hypothesised that a putative helitron-like transposable element in the large intron of *CsFT1a* may have a regulatory role in ‘FINOLA’ [8]. This putative transposable element was also found in the large intron of *CsFT1a* in ‘Boone County’ and ‘Kompolti (Figure 4c). Furthermore, considerable intron differences exist between *CsFT1b* of ‘FINOLA’, ‘Kompolti’ and ‘Boone County’ with an increase in the length of the second intron in the latter two genomes (Figure 4c), which might have functional importance in expression regulation.

The terminal flowering time (here defined as when clusters of flowers fully open at the terminal inflorescence) of ‘Boone County’ is reported as 73 and 89 days respectively for male and female individuals (Table S1). Previously ‘Kompolti’ was shown to be photoperiod-sensitive [79]. However, common accession names are not necessarily indicative of relatedness in *C. sativa* as previous work demonstrated genetic and phenotypic variation within accessions of the same name [8,90]. Nonetheless, it is unclear whether the *CsFT1* duplication can be consistently associated with photoperiod insensitivity in ‘Kompolti’ and ‘Boone County’ as has been suggested in ‘FINOLA’ [8]. It may be that the *CsFT1* (*Autoflower2*) locus interacts with another flowering locus to promote photoperiod insensitivity. In *P. sativum* and *H. vulgare* interactions among several members of the *FT* family determine photoperiodic flowering response [67,68].

## Conclusion

In conclusion, our data indicate that *PEBP* genes in *C. sativa* are of crucial importance for the reproductive transition and development and thus demonstrate significant promise to be harnessed in future breeding programmes for *C. sativa*, for example, to expand the geographic range of this crop. Based on our data, we speculate that the expansion of the *PEBP* gene family has likely been crucial in the latitudinal adaptation and evolution of sexually dimorphic flowering in *C. sativa*. This thorough characterisation of the *PEBP* family will provide the basis for the development of molecular markers for crop improvement, as well as targets for understanding the sub-functionalisation of duplicated genes.

In future work, the development of a *C. sativa* pangenome in addition to a chromosome Y assembly in *H. lupulus* will be important to address questions on the presence/absence of gene duplications and their associations with flowering phenotypes.

## Declarations

### Ethics approval and consent to participate

Not applicable.

### Consent for publication

Not applicable.

### Funding

CAD is supported by an Irish Research Council–Environmental Protection Agency Government of Ireland Postgraduate Scholarship (grant no. GOIPG/2019/1987).

### Authors’ contributions

CAD, RM, SS, PFM, and TPM designed and managed the project. CAD performed data analyses. CAD, RM, and SS wrote the manuscript. All authors revised, read, and approved the manuscript.

### Data Availability

All genome sequences are publicly available and accessible via the URLs listed in the methods. RNA seq data used is available NCBI under bioprojects PRJNA956491 and PRJNA1126191.

## Acknowledgements

CAD would like to thank Louise Ryan for helpful discussions at the commencement of this project.

## Supplementary tables

**Table S1.** *C. sativa* genomes analysed in this study.

**Table S2.** Analysis of blast hits for *PEBP*-like genes in *H. lupulus*.

**Table S3.** Gene names and identifiers for species utilised in the analyses.

**Table S4**. Normalised coverage calculation for *MFT*-like genes in *C. sativa*.

**Table S5.** Analysis of blast hits for *PEBP*-like genes in *C. sativa* cultivar ’FINOLA’.

**Table S6.** Candidate conserved amino decides to maintain PEBP function and to differentiate between PEBP subfamilies.

## Supplementary figures

**Figure S1.** *PEBP*-like genes identified in *C. sativa* and *H. lupulus* and closely related species.

**Figure S2.** Phylogenetic tree of PEBP-like genes in *H. lupulus* and ten *C. sativa* genomes.

**Figure S3.** Phylogenetic tree of *PEBP*-like genes in *C. sativa*.

**Figure S4.** Comparing conserved FT residues that indicate flowering promoter function in multiple species.

## Notes

### Competing Interest Statement

The authors have declared no competing interest.

