## Supplementary figures for "*FT*-like genes in Cannabis and hops: sex specific expression and copy-number variation may explain flowering time variation"

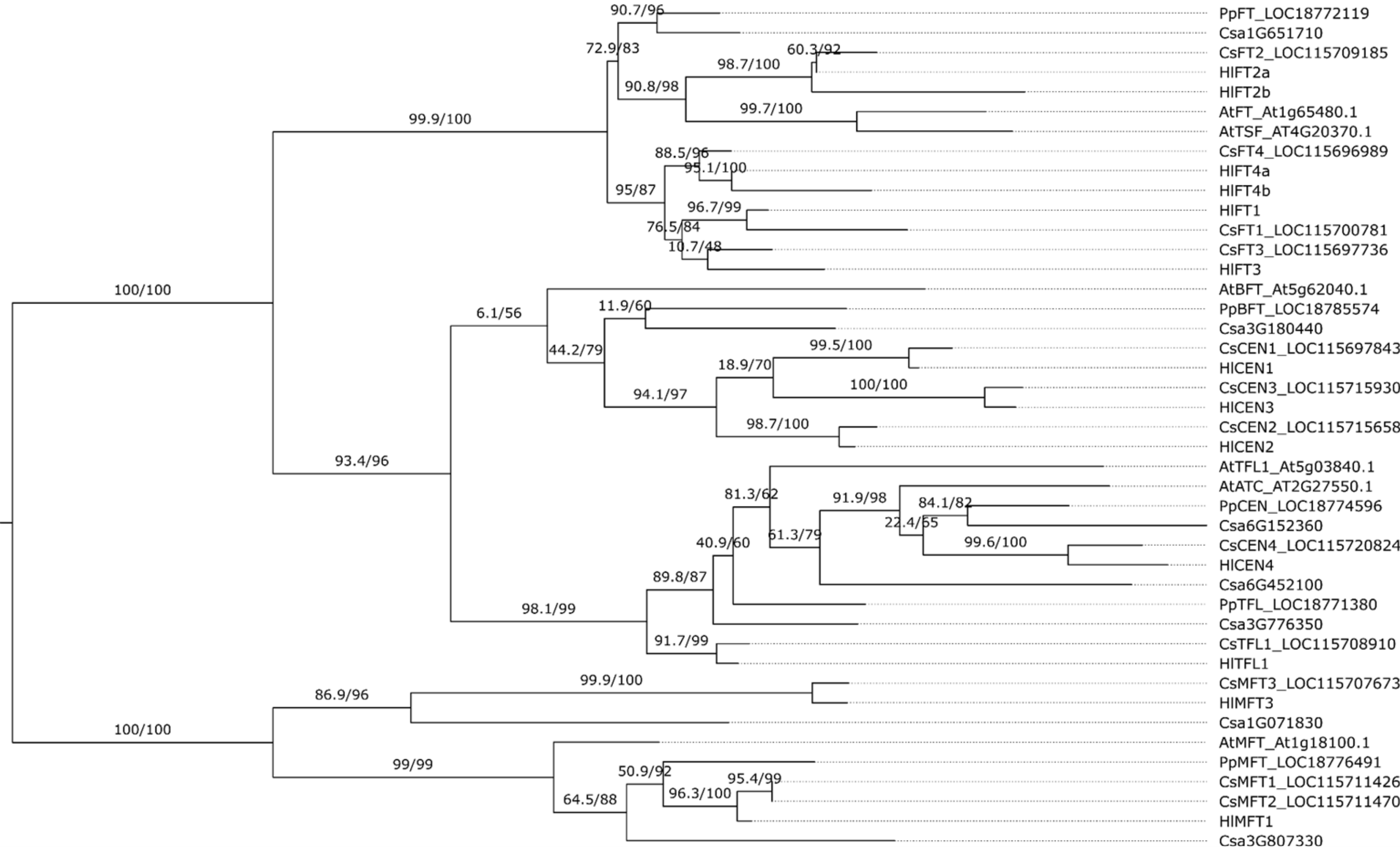


**Figure S1. *PEBP*-like genes identified in *C. sativa* and *H. lupulus* and closely related species*.*** A phylogenetic tree of PEPB-like proteins from *A. thaliana, P. persica, Cucumis sativus, C. sativa* and *H. lupulus* shows three main clades: FT, MFT and TFL1/CEN-like. *Cucumis sativus* has two MFT clades, similar to the Cannabaceae.

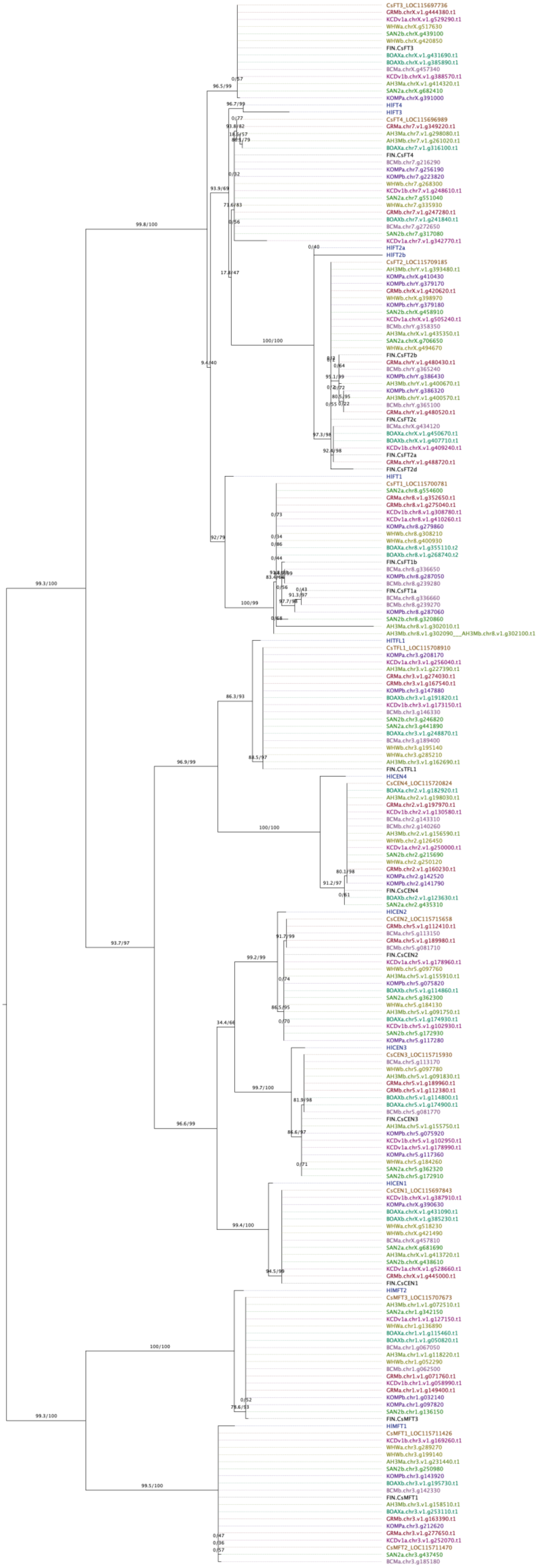

**Figure S2. Phylogenetic tree of *PEBP*-like genes in *H. lupulus* and ten *C. sativa* genomes.** An expanded version of the tree is in Figure S3. Branch support values are from Ultrafast bootstraps and a Shi-modaira–Hasegawa approximate likelihood ratio test (SH-aLRT) test (1000 replicates each).

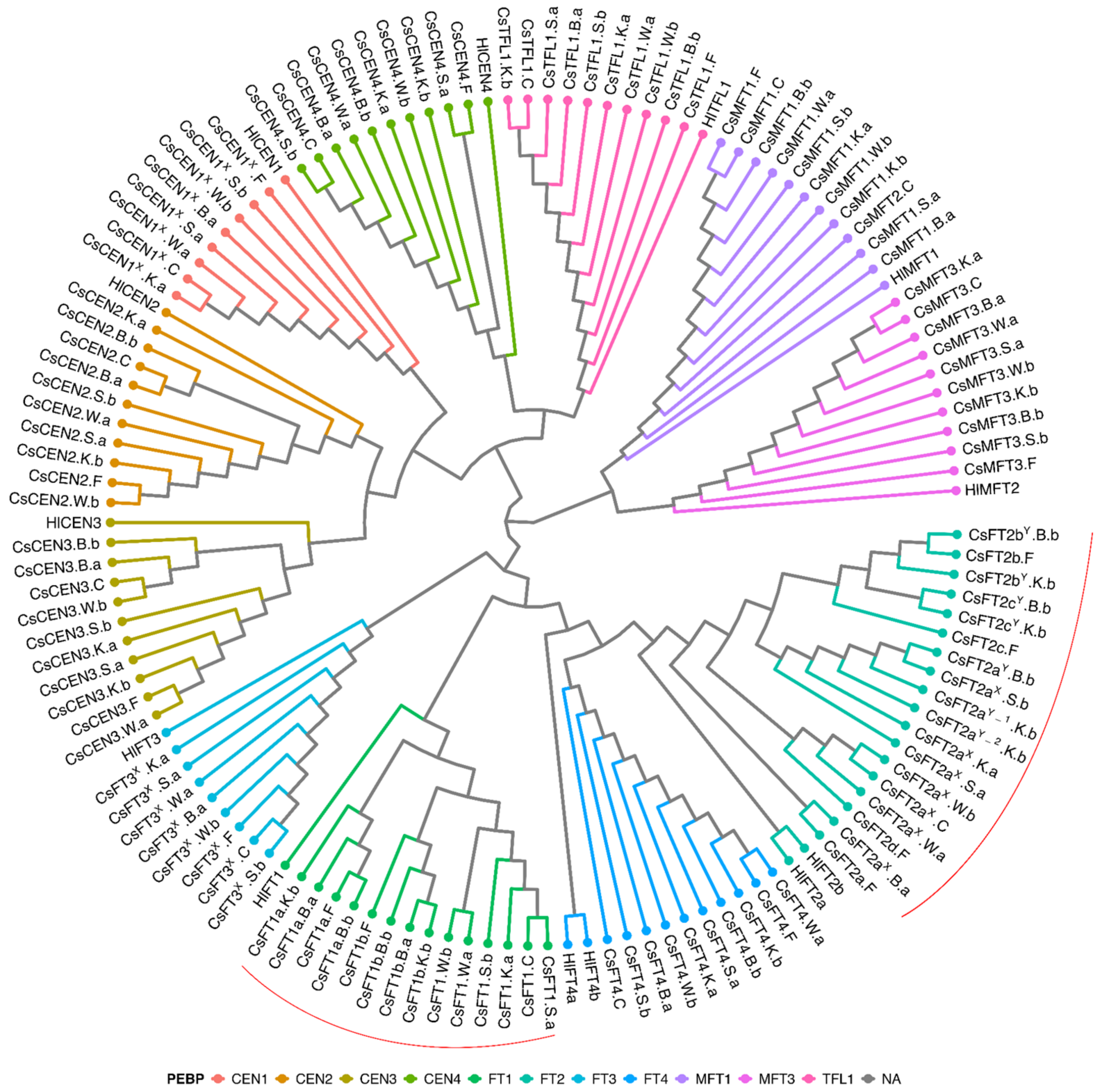


**Figure S3. Phylogenetic tree of *PEBP*-like genes in *C. sativa*.** Branch colours denote the PEBP clades. Stars highlight the gene duplications found in the *CsFT1* and *CsFT2* subclades. Tip labels include gene name (-X or -Y if located on sex chromosomes), accession (C= 'CBDRx, F= 'FINOLA', W= 'White Widow', S= 'Santhica 27', K= 'Komploti', B= 'Boone County'), and haplotype (a, b). See Figure S2 for tree with bootstrap support values. Red lines highlight gene duplications of interest.

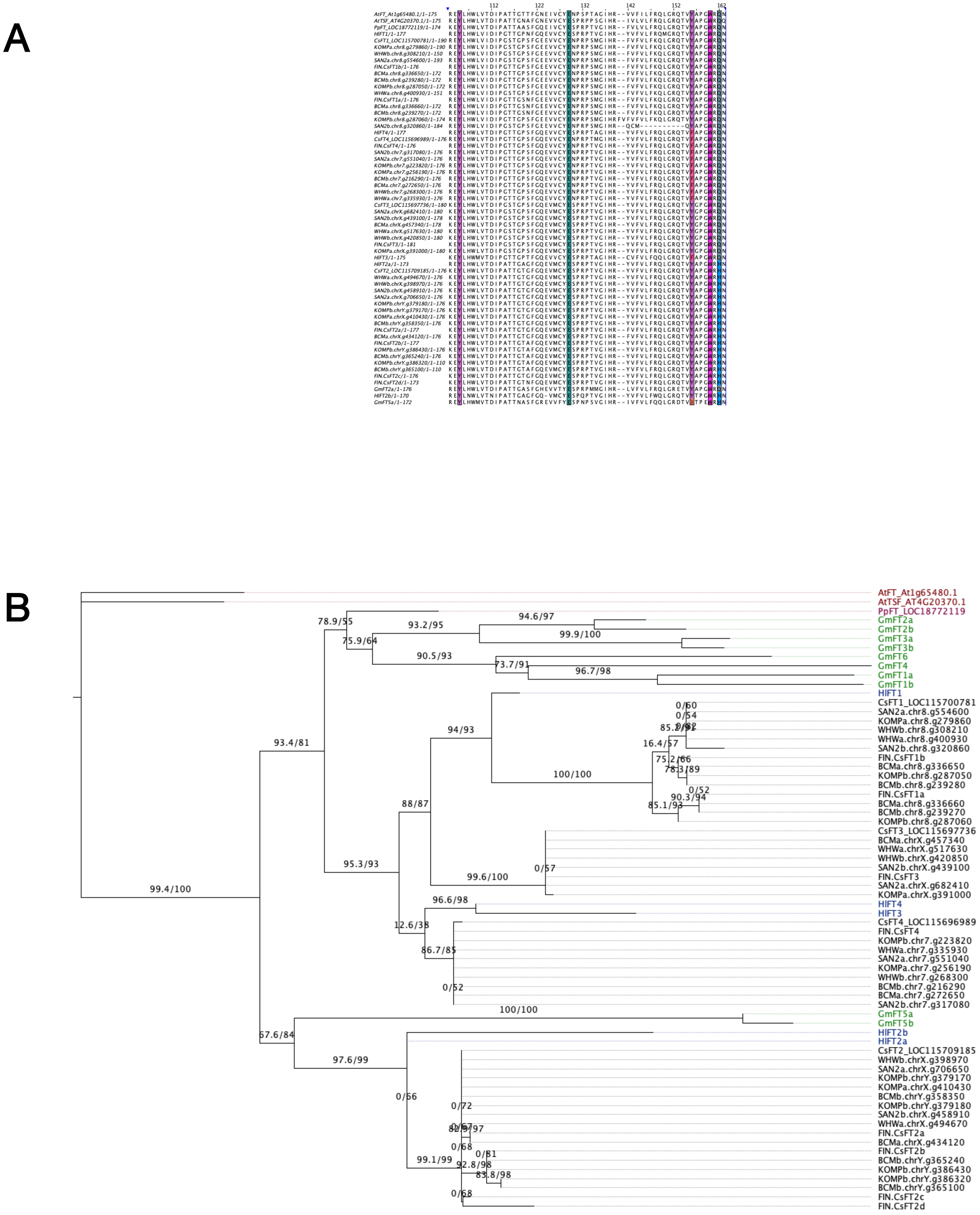


**Figure S4. Comparing conserved FT residues that indicate flowering promoter function in multiple species.** Multispecies comparison of FT-like protein sequences in *A. thaliana, P. persica, G. max, C. sativa* and *H. lupulus* (a). Five key amino acids highlighted are of known importance from the literature (position refers to AtFT): Y85, E109, Y134, W138 and Q140 (Wu *et al.*, 2017). The key amino acid positions are as expected in all *C. sativa* proteins except CsFT2 which has a substitution at Q140H. GmFT5a also has this substitution and is a demonstrated flowering promoter (Cai et al., 2020, Nan et al., 2014). There are 10 FT-like proteins in *G. max*, but for brevity, only GmFT2a and GmFT5a were included in protein alignment. The relationship between HlFT2, CsFT2 and GmFT5a was confirmed in a phylogenetic tree (b).
